# Scd1 diffuses end to end along the cytoplasm to facilitate Cdc42 activation and bipolar growth

**DOI:** 10.1101/2025.01.24.634833

**Authors:** Marcus A Harrell, Olivia Chinsen, Maitreyi Das

## Abstract

The conserved GTPase Cdc42 is a major regulator of polarized growth in most eukaryotes. In *Schizosaccharomyces pombe*, Cdc42 activity displays anticorrelated oscillatory dynamics between the growing ends enabling bipolarity. Cdc42 at each end is activated only when the opposite end loses activity. This suggests that a regulator of Cdc42 likely travels end-to-end to activate Cdc42. The oscillatory dynamics between the growing ends have also been observed in Cdc42 activator Scd1, its scaffold Scd2. It is unclear how these proteins move between the ends to facilitate bipolarity. We find that Scd1 does not travel between the cell ends via actin-mediated delivery. Instead, we show that Scd1 is mostly cytoplasmic and diffuses between the cell ends. The rate of diffusion is not entirely proportional to increasing the mass of Scd1 and cells lacking the inhibitor Pak1 kinase show decreased diffusion. Moreover, we show that Scd1 diffuses at a much faster rate compared to its scaffold Scd2. These findings suggest that Scd1 diffusion is not random and is regulated by Pak1 kinase. We find that decreasing the rate of diffusion disrupts Cdc42 oscillatory dynamics and results in monopolarity. Our results show that end-to-end Scd1 diffusion drives Cdc42 oscillatory dynamics and regulates cell polarity.

**SIGNIFICANCE STATEMENT:**

- Cdc42 activation shows oscillatory dynamics between the sites of growth
- The Cdc42 GEF Scd1 diffuses from site of activation to the opposite end to facilitate these oscillatory dynamics
- This diffusion is not random and likely depends on intrinsic properties of the Scd1 protein.

## INTRODUCTION

The Rho Family GTPase Cdc42 is a major regulator of polarized growth across most eukaryotes (Miller and Johnson, 1994; Johnson, 1999). Cdc42 is a molecular switch that is active when GTP bound, and inactivated when GTP is hydrolyzed to GDP (Johnson, 1999). Active Cdc42 promotes F-actin polymerization and membrane trafficking for cell growth. Thus, Cdc42 activity must be precisely coordinated among multiple sites of growth for proper cell polarity and function (Nobes and Hall, 1999; Stengel and Zheng, 2011; Das and Verde, 2013). GTPases are activated by GEFs (Guanine nucleotide exchange factors) and inactivated by GAPs (GTPase activating proteins) (Hall, 1990). GEFs activate Cdc42 by exchanging GDP for GTP, while GAPs complete a catalytic triad to promote hydrolysis of GTP into GDP (Zhang *et al*., 1999; Scheffzek and Ahmadian, 2005).

The fission yeast *S. pombe* has a streamlined proteome compared to more complex model systems. Cdc42 has around 20 GEFs and GAPs in mammalian cells (Rossman *et al*., 2005). But, in *S. pombe*, there are only two Cdc42 GEFs, Scd1 and Gef1, and three Cdc42 GAPs (Murray and Johnson, 2001; Coll *et al*., 2003). These streamlined signaling pathways make *S. pombe* an excellent model to investigate Cdc42 regulation. *S. pombe* has 3 major players for Cdc42 activation and growth, Gef1, Scd1, and the scaffold Scd2 (Chang *et al*., 1994; Sawin and Nurse, 1998; Chang *et al*., 1999; Wheatley and Rittinger, 2005). The initial activation of Cdc42 is mediated by Ras GTPase or Gef1 (Chang *et al*., 1994; Hercyk *et al*., 2019b). When active, GTP-bound Cdc42 recruits Scd2 (Bem1p) to tether Scd1 (Cdc24p) to the membrane and activate more Cdc42 (Butty *et al*., 2002; Wheatley and Rittinger, 2005; Rapali *et al*., 2017). This results in a positive-feedback loop wherein Cdc42 activation is self-amplified (Butty, 2002; Kozubowski *et al*., 2008; Das and Verde, 2013).

During interphase, Cdc42 activity oscillates between the two growing ends (Das *et al*., 2012). Cdc42 oscillations are anti-correlated, such that as one end gains Cdc42 activity, the other end loses activation (Das *et al*., 2012). This spatiotemporal patterning suggests that the two ends are interconnected by some protein(s) that are alternately recruited to the growing ends for oscillatory activation of Cdc42.

The identity of this factor or how it travels from one end to the other is unknown but Cdc42 GEFs are a likely candidate. Previous work has shown Gef1 and Scd1 play distinct roles in cytokinesis and Gef1 is a secondary GEF for Cdc42 activation and initiates growth at new sites but is not the major GEF needed for polarized growth (Wei *et al*., 2016; Hercyk *et al*., 2019a). Indeed, *gef1Δ* mutants are mostly monopolar, but still polarized whereas *scd1Δ* mutants are depolarized, and show spherical morphology (Hercyk *et al*., 2019a). This suggests that Scd1 is the primary GEF for cell polarity. Cdc42 oscillatory dynamics occur due to time-delayed negative feedback where the Pak1 kinase prevents Scd1 recruitment to the cell membrane (Das *et al*., 2012). We have recently shown Scd1 returns to each cell end after inhibition by Pak1 kinase in an endocytosis-dependent manner. Pak1 kinase at each end is removed during endocytosis and this allows the return of Scd1 for the next cycle of Cdc42 activation at that end (Harrell *et al*., 2024). We asked if the population of Scd1 within the cell sequentially moves from one end to the other for oscillatory Cdc42 activation.

To this end, we used *in vivo* experiments in *S. pombe* to test if Scd1 travels from one end to the other to promote bipolar growth. We find that in fission yeast, Scd1 is mostly cytoplasmic and its localization to the ends is not dependent on actin-mediated delivery. Our data shows that Scd1 diffuses between the two halves of the cell. Furthermore, Pak1 kinase promotes faster diffusion of Scd1 within the cell. We show that Scd2 also localizes via cytoplasmic diffusion, yet Scd1 molecules have a faster turnover between the two halves of the cell and their growing ends when compared to Scd2 despite Scd2 being a smaller and more abundant protein. Our data also show that simply increasing the mass of Scd1 does not linearly decrease its diffusion dynamics even though it impacts its ability to promote bipolarity. Cells with slower Scd1 diffusion show disrupted Cdc42 oscillatory dynamics and increased monopolarity. Together, our findings here show that Scd1 diffuses from one end of the cell to the other to promote oscillatory behavior but this does not occur via simple random diffusion and likely depends on the properties and regulation of Scd1.

## RESULTS & DISCUSSION

### Scd1 is highly cytoplasmic and diffuses between cell ends

It is unknown how Scd1 is delivered to the cell ends for activation of Cdc42. We asked if Scd1 localizes to the cell ends via membrane trafficking. We and others previously reported that Scd1 localization is diminished at cell ends upon depolymerization of all F-actin during LatA treatment (Hercyk *et al*., 2019a; Salat-Canela *et al*., 2021). Thus, we asked if Scd1 was delivered via actin cables. We used *for3Δ* mutants, which cannot polymerize linear actin cables for actin-mediated delivery during interphase (Feierbach and Chang, 2001; Nakano *et al*., 2002; Martin and Chang, 2006). We compared Scd1-mNG localization in *for3^+^* cells and *for3Δ* mutants and found that Scd1 localization is not decreased by the lack of actin-mediated delivery (Fig. 1A, B). Together, these data suggest that Scd1 is not localized by actin-mediated membrane trafficking. Furthermore, bipolar *for3Δ* mutants show increased Scd1 localization and anticorrelated Cdc42 oscillations thus suggesting that actin does not facilitate delivery of Scd1.

**Figure 1.**
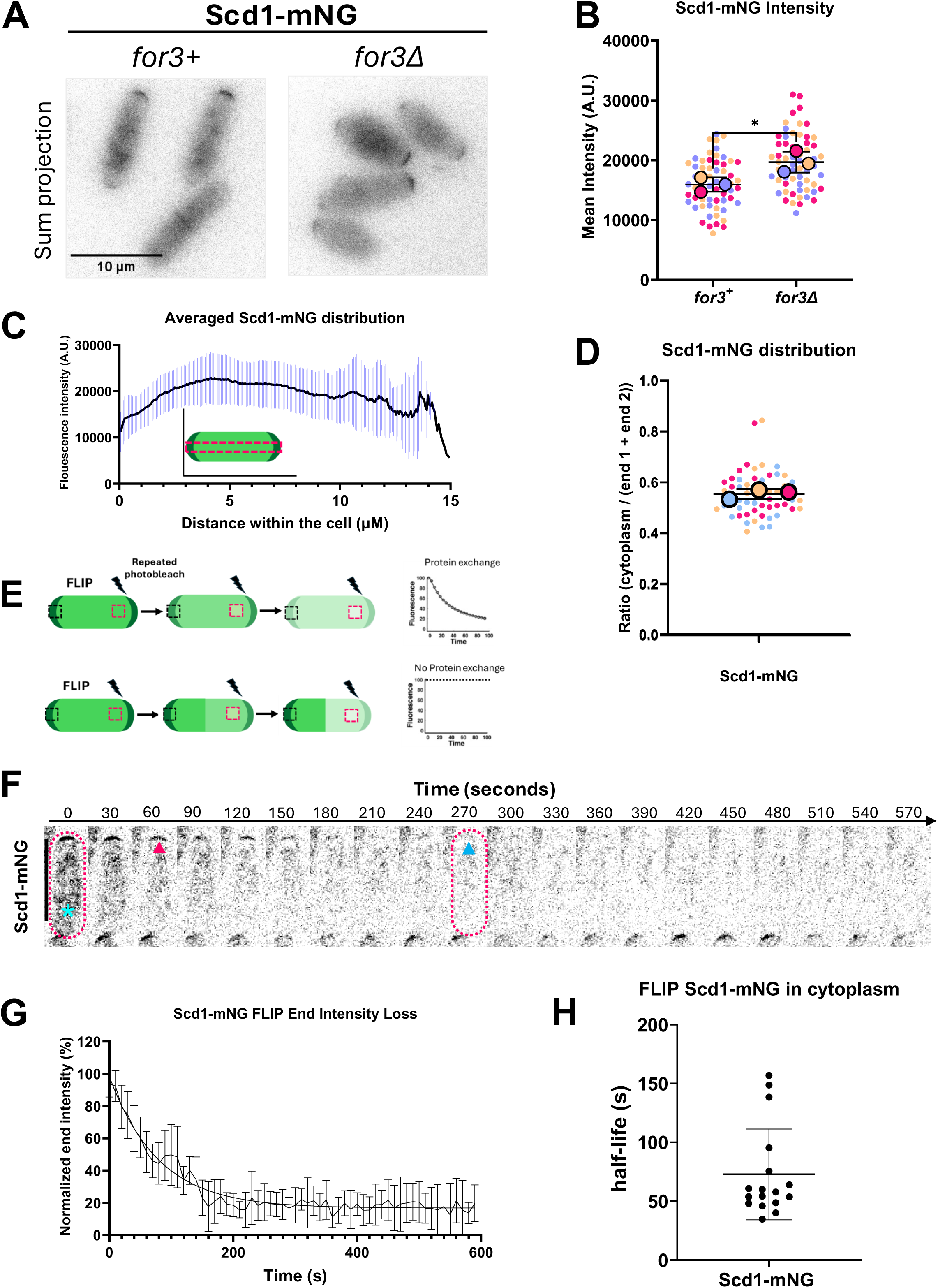
Scd1 is highly cytoplasmic and diffuses along the cytoplasm between cell ends. (A) Scd1-mNG localization in *for3^+^* cells compared to *for3Δ* mutants. Magenta rectangle indicates the cell example shown in C. (B) Quantification of mean Scd1-mNG intensity at cell ends in the indicated strains. (C) Averaged plot profile of end-to-end Scd1-mNG distribution along the long axis of the cell in G2 cells (n=59). (D) The ratio of cytoplasmic intensity relative to intensity at the cell ends (N=3, n=59). (E) Diagram illustrating fluorescence loss in photobleaching (FLIP). Top: If there is exchange of protein between the cytoplasm and the opposite cell end, we will see loss in fluorescence at the end. The fluorescence decay graphs will determine the rate of diffusion. Bottom: If there is no exchange of protein between the two halves of the cell, we will not see loss of fluorescence at the opposite cell end. (F) FLIP performed on Scd1-mNG expressing cell. Magenta dotted outline indicates the bleached cell. Cyan asterisk shows the repeatedly bleached region. Magenta arrow indicates when about 50% of the fluorescence is lost at the cell end. Blue arrow indicates roughly when all signal is lost at the cell end. (G) FLIP experiments are pooled and fitted to a one phase decay curve. (H) Quantification of the time to lose 50% of fluorescence (half-life) of Scd1-mNG in FLIP experiments. Scale bar, 10 µm. n.s., not significant; P value, *<0.05, Student’s t test.

Many yeast proteins localize in the cytoplasm as part of their function (Matsuyama *et al*., 2006). Fluorescent microscopy of Scd1 shows high cytoplasmic fluorescence intensity relative to the cell ends (Fig. 1A). We quantified the distribution of Scd1-mNG intensity throughout the cell and find that Scd1 localization within the cytoplasm is high even when compared to the more intense cell end (Fig. 1C). To quantify this metric, we averaged the intensity of the cytoplasm and compared the cytoplasmic intensity to the two cell ends and found that the cytoplasmic intensity of Scd1-mNG is about 50% of the total Scd1-mNG localized at the cell ends (Fig. 1D). This suggests that there is a high ratio of Scd1-mNG that is localized in the cytoplasm of the cell despite the low abundance of Scd1.

It has been established that the diffusion of solutes and macromolecules is important for several cellular processes (Verkman, 2002). Due to high levels of Scd1-mNG in the cytoplasm and Scd1-mNG localization occurring independent of membrane trafficking, we hypothesized that Scd1-mNG localizes to cell ends via diffusion. To test this, we used fluorescence loss in photobleaching (FLIP) (Cole *et al*., 1996; White and Stelzer, 1999; Shcheprova *et al*., 2008). FLIP is a photomanipulation technique that measures the rate of protein exchange between two regions of interest within a cell by repeatedly bleaching one area of the cell while measuring the rate of fluorescence loss at a second region of interest within the cell. If fluorescence is lost at the second region, this suggests that the two regions of interest exchange proteins (White and Stelzer, 1999). Further, the rate of this loss can be used to approximate the rate of protein exchange between these regions (Fig. 1E, top). If there is no exchange between the two regions of interest, no loss in fluorescence will be observable at the second site (Fig. 1E, bottom).

We used FLIP to test if Scd1 is exchanged between the two ends of bipolar cells or if each end has a distinct pool of Scd1. We bleached within the cytoplasm to ensure consistent photobleaching for each experiment and fluorescence loss was observed at the cell end of the opposite half of the cell from where cytoplasmic beaching occurs (Fig. 1F). We performed FLIP on Scd1-mNG cells and found that the two halves of the cell do exchange Scd1-mNG, as one half’s cytoplasm is bleached, the growing end of the opposite half steadily loses fluorescence above the rate of photobleaching (Fig. 1F and Fig. S2A). Notably, the low signal-to-noise ratio of Scd1-mNG combined with photobleaching correction, results in a plateau of intensity when all fluorescence has been lost (Fig. 1F, G). We found that the average time to lose 50% fluorescence intensity from the opposing cell end was around 75 seconds (Fig. 1H). These findings suggest that Scd1 is exchanged between the two halves of the cell and that Scd1 may localize via diffusion.

### Increasing mass on Scd1 dampens Cdc42 activation and increases monopolarity

We next asked if Scd1 dynamics can be dampened by increasing the mass on the protein. Because Scd1 is the major activator of Cdc42, dampened Scd1 dynamics likely disrupts proper Cdc42 activity. To test this, we observed CRIB-mCherry dynamics, which specifically localize to the active form of Cdc42, in Scd1-mNG and Scd1-3xGFP strains (Fig. 2A). Scd1 is a 99.1kDa protein and each fluorophore adds roughly 26.6 kDa of mass. We tested Cdc42 activation dynamics in-mNG (+26.6 kDa) and-3xGFP strains (+79.8 kDa). Indeed, we find that CRIB-mCherry oscillates between the two ends as expected in Scd1-mNG cells however CRIB-mCherry dynamics are notably more monopolar in Scd1-3xGFP cells (Fig. 2A, B). To quantify this difference, we measured how much CRIB-mCherry fluctuated between each time point at the weaker cell ends and find that Cdc42 activation is less dynamic when Scd1 is massive as seen with CRIB-mCherry in Scd1-3xGFP expressing cells (Fig. 2C).

**Figure 2.**
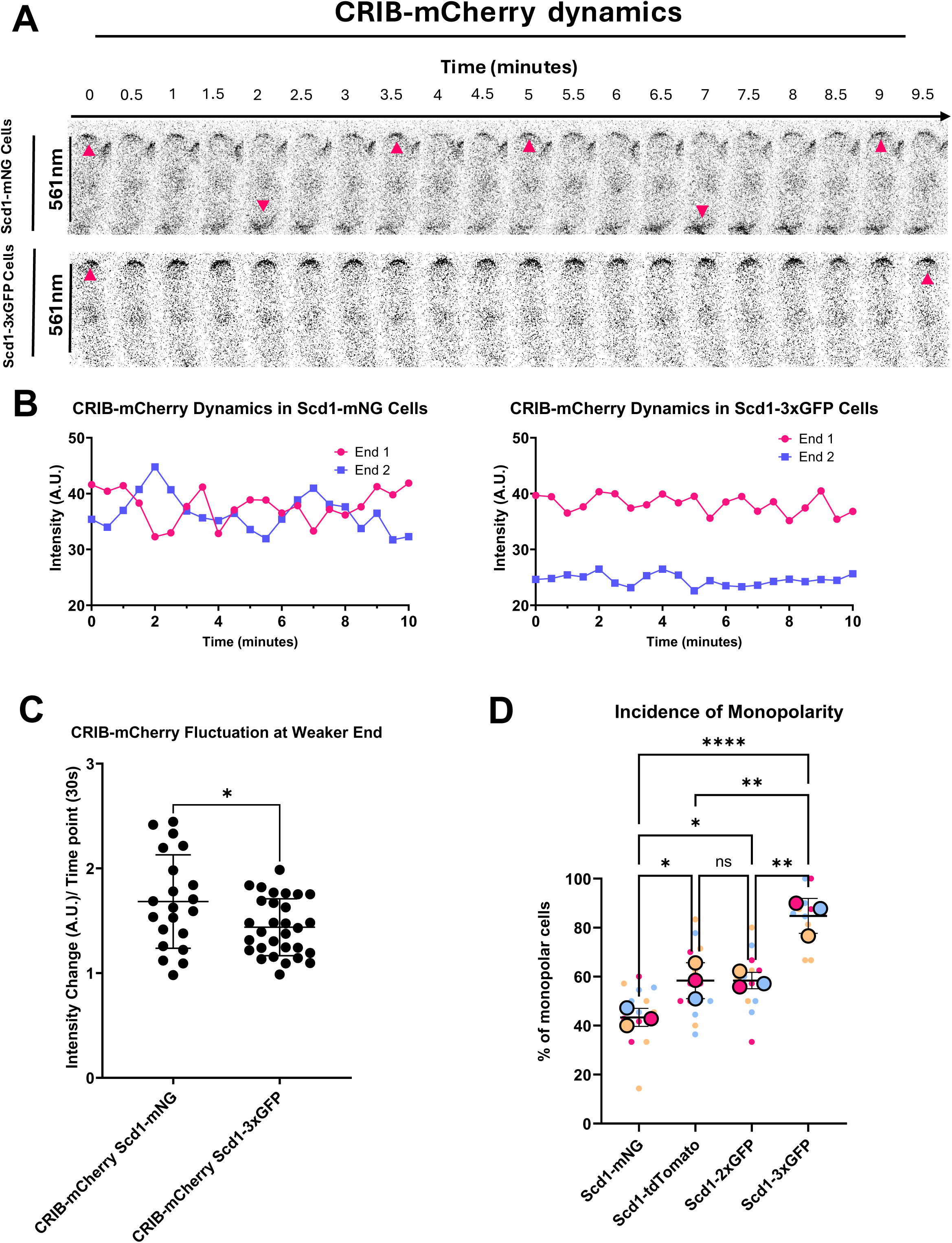
Dampened Scd1 dynamics correlate with dampened Cdc42 activation. **(A)** Montage of CRIB-mCherry dynamics in Scd1-mNG cells and Scd1-3xGFP cells. **(B)** Representative graphs of CRIB-mCherry dynamics in Scd1-mNG cells and Scd1-3xGFP cells. (C) Quantification of the extent of Pak1-mEGFP fluctuations at the cell ends (N = 3, n ≥ 5 cells). (D) Quantification of the percentage of monopolar cells observed in each culture. (N=3, n ≥ 10 cells). Scale bar, 10 µm. P value, *<0.05, **<0.005, ****<0.0001, Student’s t test and one-way ANOVA followed by Tukey’s multiple comparison test.

To test the impact of adding additional mass to Scd1 on cell polarity, we stained cells with calcofluor white to quantify the percentage of monopolar cells in cultures of cells with fluorescently tagged Scd1. In addition to-mNG and-3xGFP, we also tested strains with - tdTomato and-2xGFP (+54.2 kDa). We find there is an increased percentage of monopolar cells as more mass is added to Scd1 suggesting that increased Scd1 mass dampens bipolar Cdc4 activation and results in monopolar cells (Fig. 2D and Fig. S2A). In cells with increased Scd1 mass a smaller but consistent fraction of the cells are bipolar, suggesting that it is rare but still possible for Scd1 to accumulate at the opposite end. But under these conditions, this evolutionary important system of bipolar growth no longer consistently functions within the population (Fig. 2D).

### Increasing the mass on Scd1 does not proportionally decrease its diffusion dynamics

Since Scd1 diffuses from one end of the cell to the other and increasing the mass of Scd1 leads to increased monopolarity and altered Cdc42 activation patterns, we asked if Scd1 is distributed between ends via random diffusion in the cytoplasm. If this is true, the decrease in Scd1 diffusion should be proportionally inverse to the increase in mass of this protein. We aimed to test if Scd1 diffusion and localization to cell ends is slowed by adding more mass to Scd1. We used FLIP to test if Scd1 diffusion was slower in populations that have more mass added to Scd1 (Fig. 3A and Fig. S3A). We tested the rate of Scd1 loss in-mNG (+26.6 kDa),-2xGFP (+53.2 kDa), and-3xGFP strains (+79.8 kDa) (Fig. 3B and Fig. S3A). We find that there is a significant dampening of the rate of Scd1 loss in-2xGFP and-3xGFP strains when compared to-mNG (Fig. 3C and Fig. S3A). While the half-life of Scd1-mNG fluorescence at the cell end is still lost around 70 seconds, Scd1-2xGFP and Scd1-3xGFP cells show a half-life of around 100 seconds. This suggests that additional mass does hinder Scd1 mobility and dampens its dynamics. However, we did not see any change in the diffusion dynamics between Scd1-2xGFP and Scd1-3xGFP. This indicates that in addition to mass other factors also regulate Scd1 diffusion dynamics.

**Figure 3.**
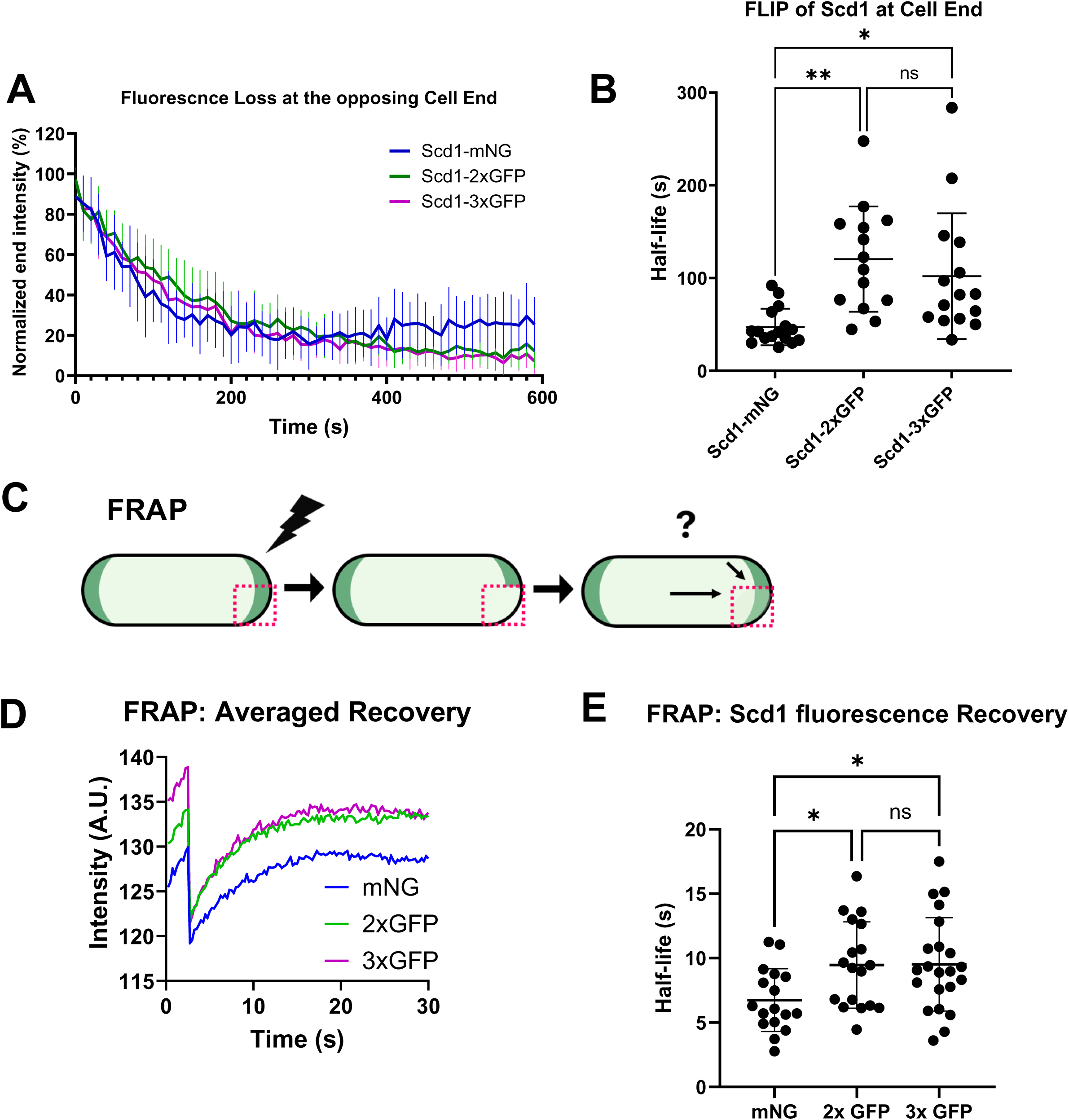
FLIP and FRAP analysis of Scd1 with additional mass. **(A)** Scd1-mNG,-2xGFP, and-3xGFP cells were subjected to FLIP and loss in fluorescence was pooled and fitted to a one-phase decay curve (n≥15). **(B)** Comparison of the half-lives of fluorescence loss in Scd1-mNG,-2xGFP, and-3xGFP cells (n≥15). **(C)** FRAP will be used to completely bleach half of one cell end and we will quantify the rate of fluorescence recovery at this cell end. (**D)** Averaged FRAP graphs for Scd1-mNG,-2xGFP and-3xGFP. (n≥13). **(E)** Quantification of the half-lives of FRAP in cells expressing Scd1-mNG,-2xGFP, and-3xGFP. (n≥17). Scale bar, 10 µm. n.s., not significant; P value, *<0.05, **<0.005, one-way ANOVA followed by Tukey’s multiple comparison test.

Next, we tested if Scd1 recruitment at the cell ends is perturbed by additional mass. To this end, we used fluorescence recovery after photobleaching (FRAP) (Poo and Cone, 1974; Axelrod *et al*., 1976; Jacobson *et al*., 1976). We bleached half of one cell end to standardize the amount of bleaching and observe if the arrival of unbleached, new protein occurred via lateral membrane diffusion or cytoplasmic diffusion (Fig. 3C). We observe that Scd1 recovery at the photobleached region occurs without diminishing the intensity of the other half of the same cell end which suggests that recovery occurs via the arrival of protein from the cytoplasm and not due to the diffusion of the protein from one side of the end to the bleached side (Fig. S3B, and Fig. S3C). Indeed, we found that Scd1 fluorescence recovery is significantly slower in - 2xGFP and-3xGFP strains when compared to Scd1-mNG (Fig. 3D, E). However, similar to our FLIP data, we did not see any difference in the FRAP recovery of Scd1-2xGFP and Scd1-3xGFP. This further suggests that Scd1 dynamics at the cell ends are only partially dependent on its mass.

### Pak1 kinase facilitates Scd1 diffusion dynamics within the cell

In budding yeast, Cdc42 activates the PAK Kinase, Cla4 (Pak1), to phosphorylate and inhibit Cdc24 (Scd1) localization at cell ends via negative feedback (Gulli *et al*., 2000; Howell *et al*., 2012; Wu and Lew, 2013; Kuo *et al*., 2014; Wu *et al*., 2015; Rapali *et al*., 2017). In fission yeast *pak1-ts* hypomorphic mutants, high amounts of Scd1 and active Cdc42 accumulate at only one end causing the characteristic monopolar and wide phenotype (Verde *et al*., 1998; Das *et al*., 2012). We asked if Pak1 kinase also regulated Scd1 diffusion within the cytoplasm. To this end, we investigated the rate of Scd1-mNG recruitment and diffusion using FRAP and FLIP in a *pak1-ts* mutant background. We used FRAP to measure Scd1-mNG recruitment to cell ends in *pak1-ts* mutants and found there is no significant difference between *pak1^+^*cells and *pak1-ts* mutants (Fig. 4A, B, C). This suggests that the lack of Pak1 kinase on Scd1 does not significantly impact Scd1-mNG recruitment to cell ends (Fig. 4A, B, C). Next, we used FLIP and measured fluorescence loss at the cell ends to quantify if the lack of Pak1 kinase impacted Scd1 diffusion. We find that Scd1-mNG intensity loss is significantly slower in *pak1-ts* mutants when compared to *pak1^+^* cells (Fig. 4D). The half-life of fluorescence loss of Scd1-mNG in *pak1^+^* cells is 74 seconds while that in *pak1-ts* mutants is 140 seconds (Fig. 4E). These data suggest that in the absence of Pak1 kinase activity, Scd1-mNG diffusion is significantly delayed and even slower than Scd1-2xGFP and Scd1-3xGFP. The mechanism of how Pak1-dependent phosphorylation of Scd1 diffusion will be investigated further in future studies.

**Figure 4.**
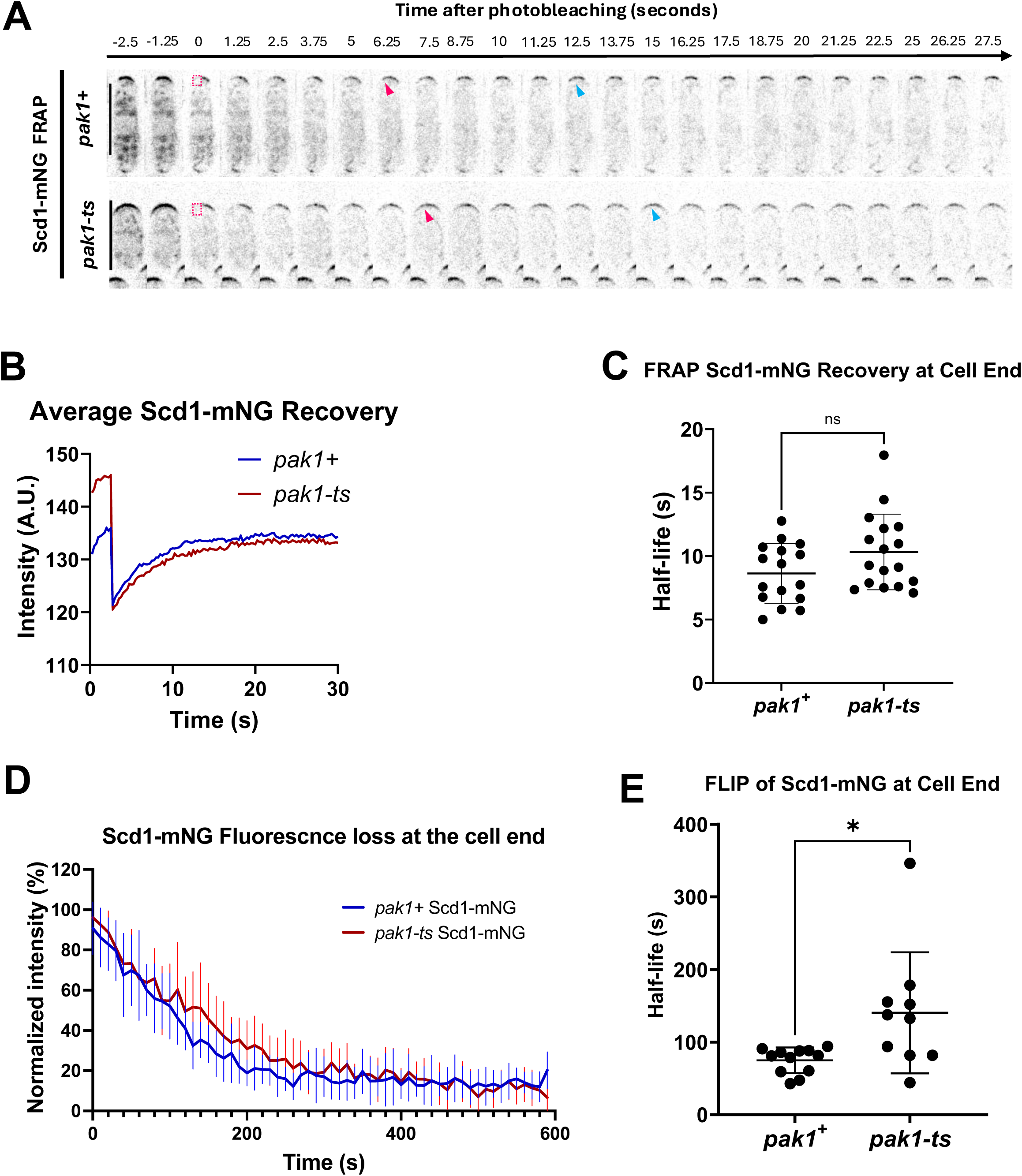
**Loss of *pak1* kinase decreases Scd1 diffusion rates. (**A) Montage of Scd1-mNG FRAP in *pak1^+^* and *pak1-ts* cells. Magenta boxes indicate the half of the cell end that has been photobleached. Magenta arrows show when half of the final fluorescence has recovered. Blue arrows indicate when the fluorescence recovery plateaus. (B) Averaged fluorescence recovery graphs of Scd1-mNG FRAP in *pak1^+^* and *pak1-ts* cells. (C) Half-life of fluorescence recovery in *pak1^+^* and *pak1-ts* cells (n≥16). (D) FLIP of Scd1-mNG in *pak1^+^* and *pak1-ts* cells cells. The fluorescence loss at the cell ends was pooled and fitted to a one-phase decay (n≥10). (E) Quantification of Scd1-mNG loss at the cell ends of *pak1^+^*and *pak1-ts* cells (n≥10). Scale bar, 10 µm. n.s., not significant; P value, *<0.05, Student’s t test.

### The scaffold Scd2 shows slower diffusion dynamics compared to Scd1

We asked if like Scd1, its scaffold Scd2 also diffused from one end of the cell to the other with similar dynamics? It is possible that the dynamics of Scd2, the Scd1 scaffold, dictates the rate of Scd1 diffusion to the sites of growth. To test this possibility, we compared the rates of diffusion between Scd1-mNG and Scd2-GFP. We began by comparing the distribution of Scd1-mNG within the cell to Scd2-GFP (Fig. 5A, B). We again see that Scd1-mNG has roughly 50% of its intensity in the cytoplasm when compared to the cell ends. By contrast, Scd2-GFP only shows about 25% of its intensity in the cytoplasm relative to the Scd2 intensity at the cell ends (Fig. 5A, B, C). Moreover. While we see a consistent dip in Scd1 levels in the cytoplasm right next to the end with increased levels, we do not such a dip for Scd2 near the ends (Fig. 5B, red arrow).

**Figure 5.**
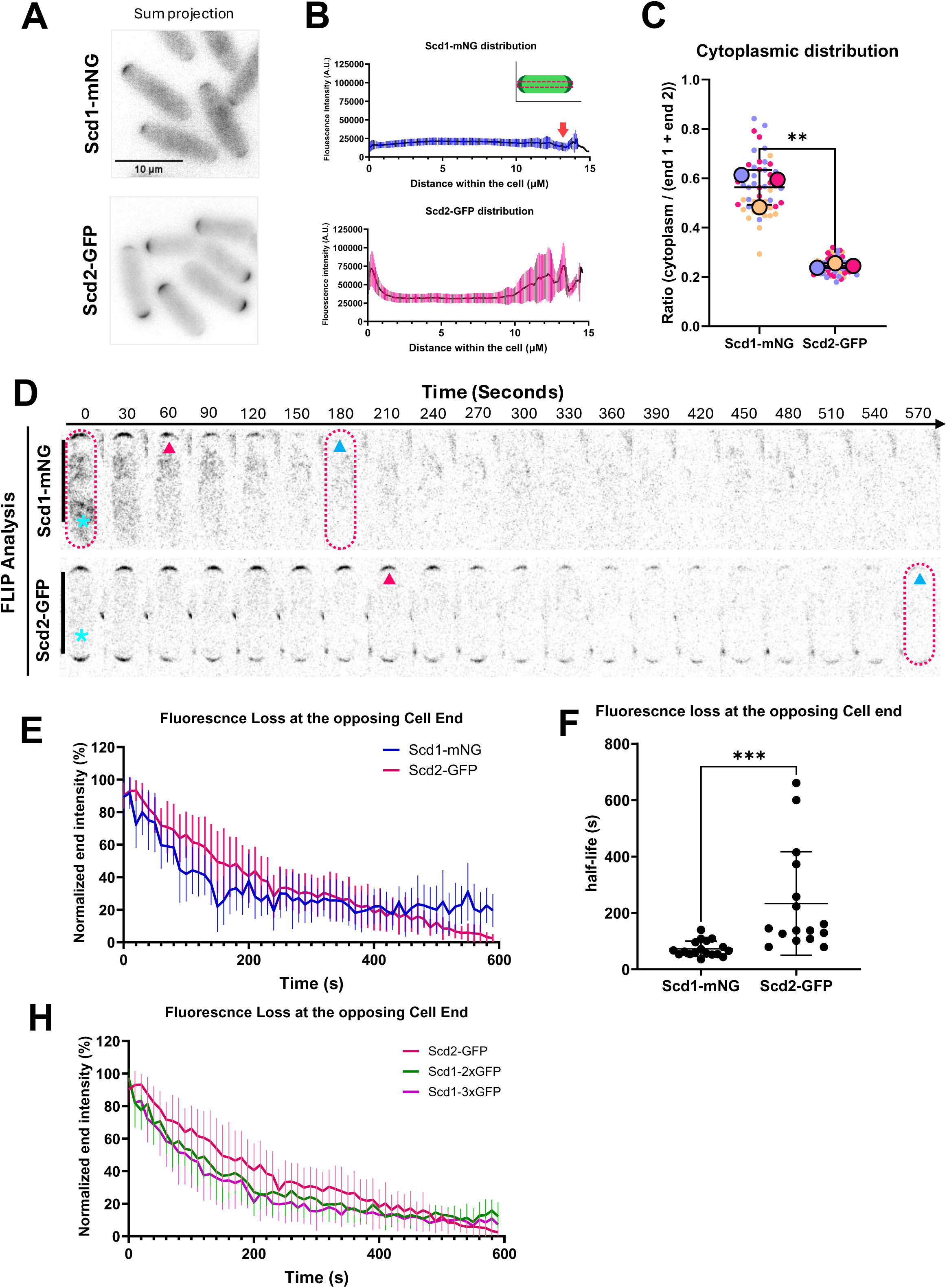
FLIP reveals that Scd1 diffuses between the cytoplasm and cell ends more rapidly than Scd2. (A) Cellular localization of Scd1-mNG and Scd2-GFP. (B) End-to-end fluorescence intensity profiles of Scd1-mNG and Scd2-GFP along the long axis of the cell (n= 60). (C) Ratio of cytoplasmic intensity relative to the intensity at the cell ends for Scd1-mNG compared to Scd2-GFP (N=3, n = 20 cells per replicate). (D) Montages of FLIP in Scd1-mNG and Scd2-GFP. Magenta dotted line indicates the bleached cell. Cyan asterisk shows the repeatedly bleached region. Magenta arrow indicates roughly when 50% of fluorescence is lost at the cell end. Blue arrow indicates roughly when all signal is lost at the cell end. (E) Pooled fluorescence loss for Scd1-mNG and Scd2-GFP fitted to a one phase decay curve (n ≥16). (F) Half-lives of fluorescence loss in Scd1-mNG compared to Scd2-GFP (n ≥16). (H) One phase decay curve of fluorescence loss of Scd2-GFP data repurposed from Figure 4 and Scd1-2xGFP and-3xGFP data repurposed from Figure 2 (n≥15). Scale bar, 10 µm. n.s., not significant; P value, **<0.005, ***<0.0009, Student’s t test.

Next, we performed FLIP on Scd1-mNG cells and Scd2-GFP cells to compare their rates of exchange between the two halves of the cell (Fig. 5D). We find that like Scd1-mNG, Scd2-GFP is also exchanged between the cytoplasm and cell ends. However, Scd1-mNG intensity is rapidly lost from the cell end whereas Scd2-GFP loss is slower (Fig. 5D, E). The average half-life of fluorescence intensity loss for Scd1-mNG is around 70 seconds on average whereas the half-life of Scd2 loss is over 200 seconds (Fig. 5E, F). Together, these data suggest that while Scd1 diffuses from one cell end to the other for bipolar Cdc42 activation, Scd2 diffusion is much slower and thus likely not a factor in Cdc42 oscillatory dynamics. Indeed, the fluorescence decay of Scd2-GFP at the opposite end is even slower than that Scd1-2xGFP and-3xGFP (Fig.5H). Thus, Scd1 (99 kDa) has a higher ratio of cytoplasmic localization and is exchanged rapidly between growing ends despite Scd2 (60 kDa) being the smaller and more abundant protein. This further suggests mass is not the only factor that impacts diffusion and the intrinsic properties of Scd1 facilitate its rapid diffusion.

### Conclusions

We previously found that Pak1 removal via endocytosis is vital for Scd1 localization and allows for the repeated cycle of Cdc42 activation at a cell end (Harrell *et al*., 2024). Here we show that Scd1 localizes to the growing cell ends via diffusion between those ends. Thus, Pak1 phosphorylation must facilitate adequate Scd1 turnover at cell ends such that there is enough cytoplasmic Scd1 molecules to diffuse to the other end to drive Cdc42 activation between the different sites of growth. Thus, we find yet another layer of self-regulation in this polarity system where Scd1 diffusion within the cytoplasm activates Cdc42 and promotes bipolar growth.

FLIP analysis of Scd1-3xGFP shows that when the half-life is slowed to around 100 seconds or more, most cells in the population are monopolar. This suggests Scd2 is exchanged less rapidly while Scd1, with a half-life of 70 seconds, must be rapidly exchanged to promote proper competition and bipolar growth. While we see differences in the diffusion rate of Scd1 and Scd2, its rate of recruitment at the cell ends as determined by FRAP is similar (Harrell *et al*., 2024). It is possible the once Scd1 is available at an end via diffusion, local Scd2 at that end facilitates Scd1 recruitment to the plasma membrane. In such a scenario, Scd2 diffusion between ends would not be necessary for Cdc42 oscillatory dynamics. The finding that Scd2 diffusion is slower than the diffusion of Scd1 suggests that this diffusion is not random and that the intrinsic properties of the proteins impact their diffusion rates. Scd1 has a higher amount of intrinsically disordered regions when compared to Scd2. While the functional significance of intrinsically disordered regions remain to be fully understood, these regions may have roles in facilitating Scd1 diffusion, possibly by making the protein more soluble when diffusing. The intrinsically disordered regions or other properties of Scd1 may explain why simply increasing the mass beyond 54.2kDa (-2xGFP and-3xGFP) alone does not further dampen Scd1 dynamics.

There is a consistent drop in cytoplasmic Scd1 intensity near the active growing end. More investigation is needed to test the significance of this pattern. It is possible that this is a region of active recruitment and turnover, so most Scd1 molecules that are within this region are rapidly recruited to the membrane of the growing cell end. Scd1 would need to be beyond a certain threshold of an active end to have a chance of diffusing to the opposite cell end. This threshold may be easier to overcome when an end is saturated and reaches its carrying capacity such that it can no longer accumulate more Scd1.

Scd1 diffusion appears to slow down in mutants of the *pak1* kinase. In Pak1 kinase mutants, Scd1 is not removed from the cell ends via negative feedback (Das *et al*., 2012). In these cells Scd1 accumulates at only one cell end and the cells are monopolar. These mutants do show cytoplasmic Scd1 with similar recruitment rates at the growing cell end similar to that of *pak1^+^* cells. However, they display slower diffusion rates suggesting that Pak1 kinase may play a role in regulating Scd1 diffusion. Indeed, the half-life of fluorescence decay in *pak1-ts* mutants is even slower than that of Scd1-3xGFP. This further highlights how Scd1 diffusion is dependent on its intrinsic properties and not just on its mass.

Scd1 is recruited to the cell membrane in its dephosphorylated form (Das *et al*., 2012). Given that our data show that Scd1 diffuses from one end of the cell to the next, it raises the possibility that a phosphatase is involved in this regulation where it dephosphorylates Scd1, thereby enabling membrane recruitment and Cdc42 activation. Such a kinase-phosphatase based regulatory system along with diffusion in the cytoplasm can allow the periodic redistribution of Scd1 at the cell ends to promote bipolar growth. Further investigations will define if similar diffusion dependent localization of Cdc42 GEFs also occur in animal cells to enable multipolar growth.

While the role for protein diffusion in the cytoplasm has not been clearly understood in biological systems, a recent report has shown that increasing cytoplasmic viscosity impairs protein diffusion in the cell and this can impact dynamic cellular processes (Molines *et al*., 2022). Cytoplasmic viscosity impacts protein diffusion in both fission yeast cells as well as in animal cells. Thus, this may be a conserved form of regulation and further studies will define the mechanistic details of diffusion of Scd1 and other proteins in polarity control.

## MATERIALS AND METHODS

### Strains and Cell Culture

The *S. pombe* strains used in this study are listed in Table S1. All strains are isogenic to the original strain PN567. Cells were cultured in yeast extract (YE) medium and grown exponentially at 25°C unless specified otherwise. Standard techniques were used for genetic manipulation and analysis (Moreno *et al*., 1991). Cells were grown exponentially for at least three rounds of eight generations each before imaging.

### Microscopy

Imaging was performed at room temperature (23–25°C). We used a Nikon Ti2 Eclipse wide-field microscope with a 100×/1.49 NA objective and an ORCA-FusionBT digital camera (Hamamatsu Model: C15440-20UP Serial No: 500428). Images were acquired using Nikon NIS Elements (Nikon). Fluorophores were excited with an AURA Light Engine system (Lumencor).

Microscopy was also performed with a 3i spinning disk confocal using a Zeiss AxioObserver microscope with an integrated Yokogawa spinning disk (Yokogawa CSU-X1 A1 spinning disk scanner) and a 100×/1.49 NA objective. Images were acquired with a Teledyne Photometrics Prime 95b back illuminated sCMOS camera (Serial No: A20D203014). Images were acquired using SlideBook (3i Intelligent Imaging innovations).

### Acquiring and Quantifying Fluorescence Intensity

Fluorescence intensity was measured using ImageJ software. All images were sum projected and mean intensities were reported. Freehand ROIs were used to measure the signal at the cell ends. For analysis of two ends of the same cell, cytoplasm with little or no signal was used for background subtraction. For all other experiments, a region outside of the cell was used for background subtraction. The mean intensity was measured and recorded after the background subtractions.

For FLIP analysis, cells were imaged every 10 s for 10 minutes with 4% laser power at 488nm wavelength. Bleaching repeatedly occurs every 2^nd^ timepoint (1 bleach every 20 seconds). The resulting FLIP data was fitted to a one-phase decay curve to indicate the time to lose 50% of fluorescence at the opposite cell end for each cell in each experiment.

For FRAP analysis, cells were imaged every 250 ms for 30 s with 15% laser power at 488nm wavelength. Square 10 x 10 Pixel (1.1 × 1.1 μm) ROIs were used to target, bleach, and measure the fluorescence intensity at half of one cell end as it was bleached and then allowed to recover. The first nine time points were acquired before photobleaching and served as a measurement of the initial fluorescence intensity. On the 10th time point, the acquisition was halted momentarily and the region of interest was photobleached using three 4% laser power bleach repetitions with a duration of 5 ms each (15 ms total). After the region of interest was bleached, acquisition resumed at 250 ms intervals as the fluorescence intensity recovered. The half-life of fluorescence recovery was quantified using SlideBook FRAP analysis software.

To quantify signal stability at the cell ends, cells were imaged every 30 s for 20 min. The stability of the proteins was then quantified by measuring the difference in intensity between each subsequent time point for the duration of the timelapse imaging. Populations that had larger average variations of fluorescent intensity between each timepoint were more dynamic and less stable than populations that had smaller variations between each timepoint.

### Statistical tests

Statistical significance was computed with GraphPad Prism. When comparing two conditions, Student’s t-test was used (two-tailed, unequal variance). One-way ANOVA, followed by Tukey’s multiple comparison tests, was used to determine significance of experiments with three or more conditions. SuperPlots (Lord *et al*., 2020) were made using GraphPad Prism.

## Acknowledgements

We thank Bret Judson and the Boston College Imaging Core for infrastructure and support. We thank Sophie Martin, and Fulvia Verde for providing strains. This work is supported by the National Institutes of Health grant R01GM136847 to M.D.

