## Supplemental Material for "Scd1 diffuses end to end along the cytoplasm to facilitate Cdc42 activation and bipolar growth"

### Supplemental Figure 1

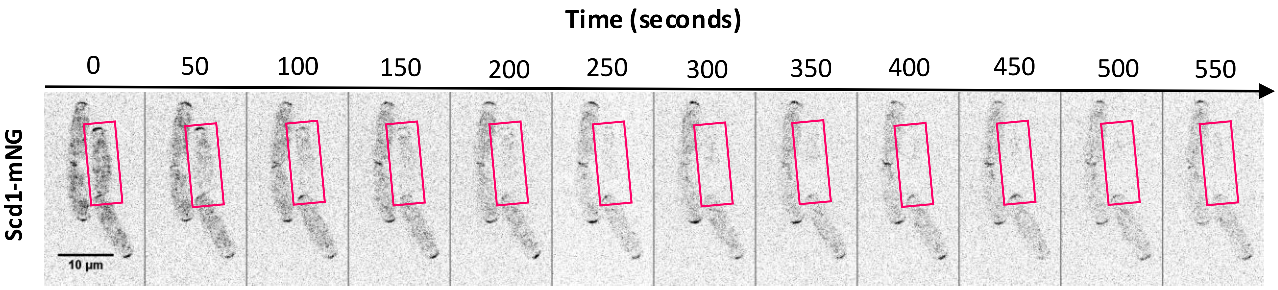


**Supplemental Figure 1. Montage of FLIP in Scd1-mNG expressing cell.** Magenta box shows the repeatedly bleached cell which is also shown in Fig. 2F

### Supplemental Figure 2

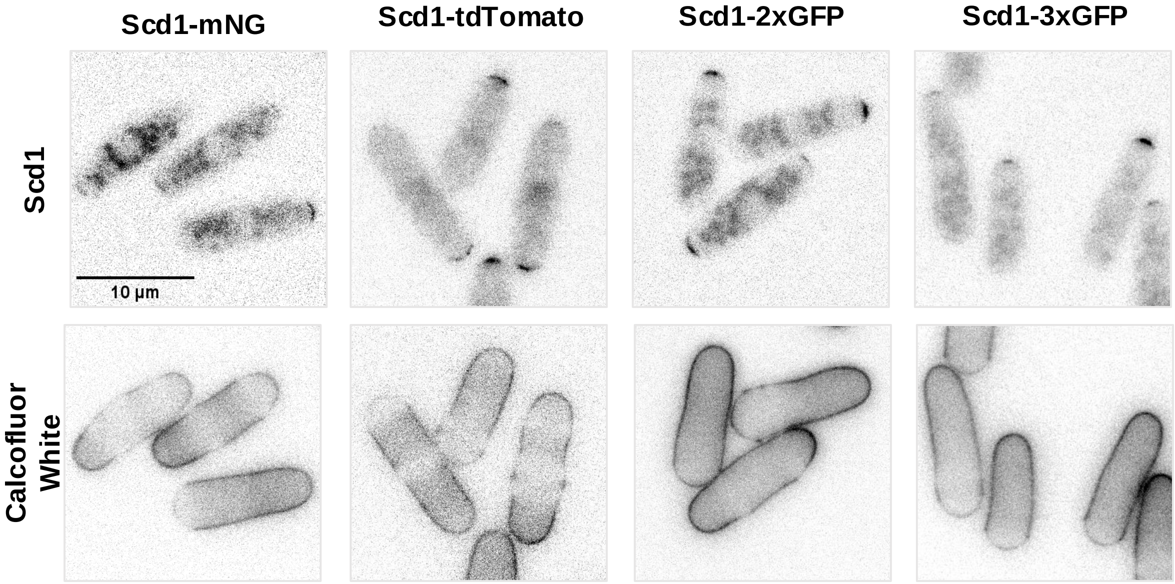


**Supplemental Figure 2. Increasing the mass of Scd1 leads to a higher incidence of monopolar cells within a population.** Fluorescence image and Calcofluor white staining of indicated strains, Scd1-mNG, Scd1-tdTomato, Scd1-2xGFP, and Scd1-3xGFP.

### Supplemental Figure 3

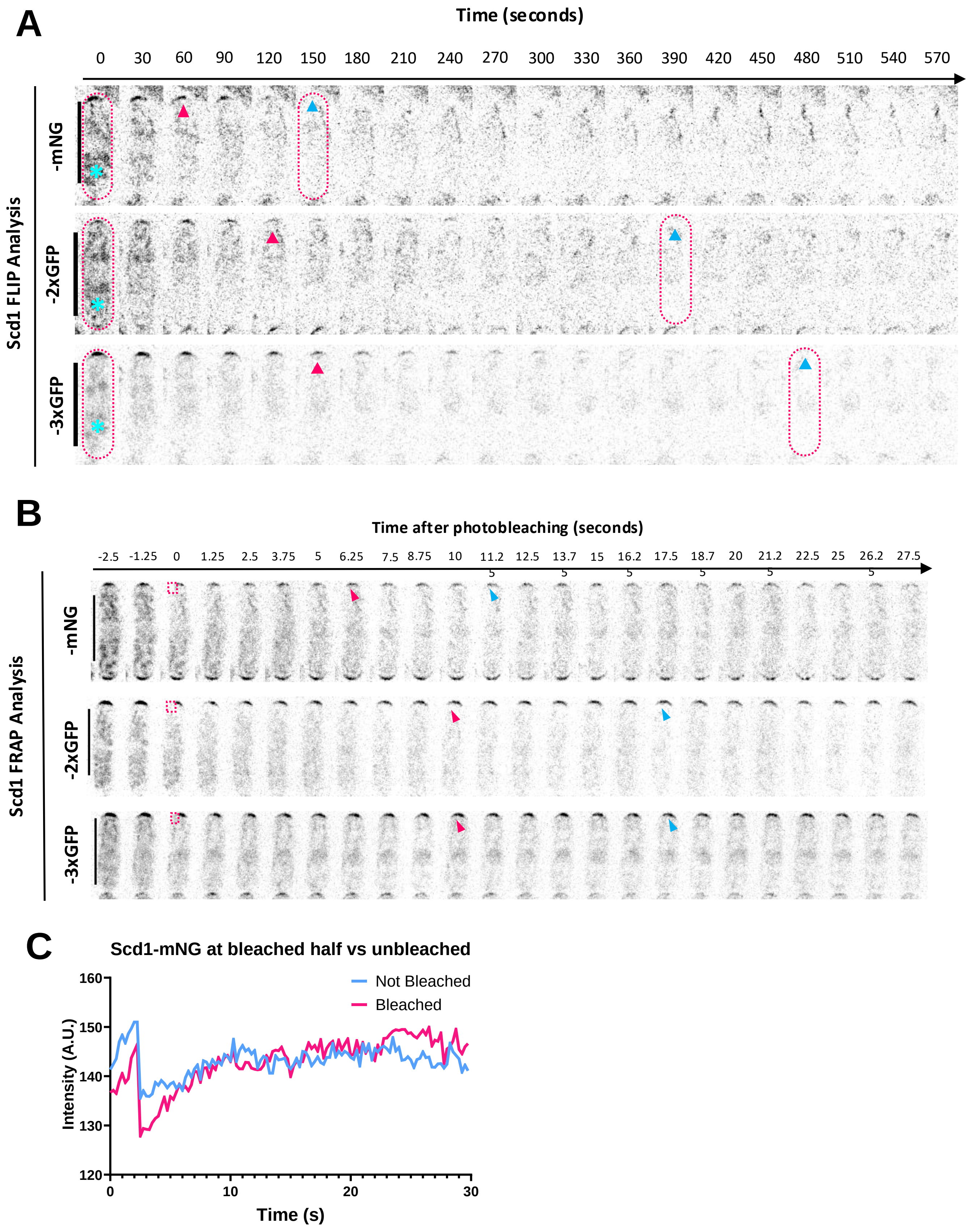

**Supplemental Figure 3. The increased mass on Scd1 decreases the rate of Scd1 diffusion and Scd1 recovery at the cell ends. (A).** Montages of fluorescence loss in Scd1-mNG, -2xGFP, and -3xGFP cells during FLIP experiments. Magenta dotted line indicates the bleached cell. Cyan asterisk shows the repeatedly bleached region. Magenta arrow indicates roughly when 50% of fluorescence is lost at the cell end. Blue arrow indicates roughly when all signal is lost at the cell end. (**B)** Montages of FRAP in cells expressing Scd1-mNG, -2xGFP, and -3xGFP. Magenta boxes indicate the half of the cell end that has been photobleached. Magenta arrows show when half of the final fluorescence has recovered. Blue arrows indicate when the fluorescence recovery plateaus. **(C)** Example graph of Scd1-mNG recovery at the bleached and non-bleached half of the cell end.

**Table S1 Strain List**

| Strain | Genotype | Origin |
|---|---|---|
| PN 567 | *h+ ade6-704 leu1-32 ura4-d18* | P. Nurse |
| FV1118 | *Scd2-GFP:KanMx ade6-704 leu1-32 ura4-d18* | F. Verde |
| FV1119 | *Scd1-GFP:KanMx ade6-704 leu1-32 ura4-d18* | F. Verde |
| YSM 947 | *Scd1-3xGFP:KanMX ade6-m216 leu1-32 ura4-d18* | S. Martin |
| YMD 1191 | *Scd1-tdTomato:KanMx ade6-704 leu1-32 ura4-d18* | Das Lab |
| YMD 1192 | *Scd1-mNG:KanMx ade6-704 leu1-32 ura4-d18* | Das Lab |
| YMD 1784 | *orb2-34 Scd1-mNG:KanMx ura4-d18 leu1-32* | Das Lab |
| YMD 1441 | *for3Δ::KanMx Scd1-mNG:KanMx leu1-32 ura4-d18* | Das Lab |
| YMD 2327 | *Scd1-3xGFP:KanMX ade6-m216 leu1-32 ura4-d18 his+* | S. Martin |
| YMD 2606 | *Scd1-mNG:KanMx CRIB-mCherry: ade-, leu1-32, ura4-d18* | This Study |
| YMD 2651 | *Scd1-3xGFP:KanMx CRIB-mCherry: ade-, leu1-32, ura4-d18* | This Study |
